# High-speed 4D fluorescence light field tomography of whole freely moving organisms

**DOI:** 10.1101/2024.09.16.609432

**Authors:** Kevin C. Zhou, Clare Cook, Archan Chakraborty, Jennifer Bagwell, Joakim Jönsson, Kyung Chul Lee, Xi Yang, Shiqi Xu, Ramana Balla, Mark Harfouche, Donald T. Fox, Michel Bagnat, Roarke Horstmeyer

## Abstract

Volumetric fluorescence imaging techniques, such as confocal, multiphoton, light sheet, and light field microscopy, have become indispensable tools across a wide range of cellular, developmental, and neurobiological applications. However, it is difficult to scale such techniques to the large 3D fields of view (FOV), volume rates, and synchronicity requirements for high-resolution 4D imaging of freely behaving organisms. Here, we present reflective Fourier light field computed tomography (ReFLeCT), a new high-speed volumetric fluorescence computational imaging technique. ReFLeCT synchronously captures entire tomograms of multiple unrestrained, unanesthetized model organisms over multi-millimeter 3D FOVs at 120 volumes per second. In particular, we applied ReFLeCT to reconstruct 4D videos of fluorescently labeled zebrafish and *Drosophila* larvae, enabling us to study their heartbeat, fin and tail motion, gaze, jaw motion, and muscle contractions with nearly isotropic 3D resolution while they are freely moving. As a novel approach for snapshot tomographic capture, ReFLeCT is a major advance towards bridging the gap between current volumetric fluorescence microscopy techniques and macroscopic behavioral imaging.

## 1 Introduction

Non-invasive tomographic 3D imaging has revolutionized basic scientific and medical research by revealing the internal structure of thick, volumetric specimens in their native biological context. However, dense tomographic 3D imaging over large fields of view (FOVs) requires a very large number of spatially resolved measurements - orders of magnitude more data than 2D imaging. It is thus challenging to efficiently capture dense tomographic image datasets at high speeds without the introduction of motion artifacts. To perform 3D tomographic imaging of whole organisms, one generally must resort to chemical fixation, immobilization, or movement restraint, which naturally disturbs the organism’s natural physiological state. An ideal tomographic imaging system would be able to synchronously capture 3D measurements of *dynamic* specimens for live 3D monitoring. Apart from avoiding the confounding effects of immobilization, such a capability would also open up new scientific opportunities for jointly observing organism behavior, morphology, and fluorescent activity, all seamlessly across 3D space and time.

Point-scanning techniques such as confocal microscopy [1] and multiphoton microscopy [2], as well as parallelized versions [3, 4], can be slow due to the need to perform inertially-constrained scanning of focused points in three dimensions. The inherent asynchrony of the acquisition of the 3D points comprising the volume of view leads to “rolling shutter” artifacts. Point-scanning techniques also tend to require high focused laser intensities, as the per-pixel integration time decreases with increasing frame or volume rate. Light sheet microscopy techniques have alleviated many of these limitations and achieved impressive volumetric imaging rates owing to its high-speed parallel 2D detection [5–9]. However, they still require mechanical scanning in one dimension, thus imposing an inverse relationship between volume rate and number of depth planes, which are acquired asynchronously. Further, at a fixed volume rate, the focused intensity of the light sheet must increase with the number of depth planes to achieve the same SNR. Light sheet and point-scanning microscopy approaches alike also typically have limited FOVs and thus often require organism immobilization.

Computational reconstruction techniques such as those used in optical projection tomography (OPT) [10] and optical diffraction tomography (ODT) [11, 12] can require sequential acquisition of hundreds of multi-angle images, which compromises speed, especially if mechanical translation or rotation is required. Multiplexing and compressive sensing can reduce the number of acquisitions and thus increase imaging speed [13, 14], albeit at the cost of reconstruction quality. Optical coherence tomography (OCT) [15, 16] is another class of wide-FOV, high-speed volumetric imaging. Even as faster sources pushing multi-MHz A-scan rates are being developed[17], OCT is also constrained by mechanical scanning speeds and lack of functional image contrast.

On the other hand, light field microscopy (LFM) and related techniques [18–23] capture 3D information synchronously in a single snapshot, requiring no moving parts and thus are limited only by the camera rate and noise. However, the acquired 3D information must be encoded onto a single 2D image sensor, which both complicates dense volumetric sampling and introduces fundamental data rate limitations. 3D techniques that perform data undersampling and use compressive sensing techniques can help address some of these shortcomings [21, 24]. While useful in certain applications, these methods often rely on strong regularizers or priors that can erase features and are not applicable when sparsity assumptions are not met. Additionally, the 3D reconstruction quality is limited by the missing cone problem due to practical design constraints of refractive objective lens, on which most LFM designs are based, that restrict the angular coverage to less than 2*π* steradians. A further complication of practical refractive objective lens designs is that achieving larger angular coverage (i.e., high numerical aperture (NA)) comes at the cost of field of view (FOV) and working distance, rendering imaging of freely behaving organisms challenging.

Here, we present a new type of high-throughput computational tomographic imaging technique based on a reflective Fourier light field design that allows for super-video-rate volumetric imaging of unrestrained organisms (e.g., freely swimming zebrafish and fruit fly larvae) over tens-of-cubic-millimeter FOVs (Fig. 1; Video 1). Key to our design is a reflective concave mirror, which overcomes many limitations of refractive objectives, enabling high angular coverage that can approach 4*π* steradians [25] with large working distances [26]. In particular, our design combines a reflective parabolic mirror objective [27] with an array of 54 individual cameras to synchronously acquire multi-view information across nearly 2*π* steradians at up to 120 Hz (multiple gigavoxels/sec). We call our method <u>R</u>eflective <u>F</u>ourier <u>L</u>ight field <u>C</u>omputed <u>T</u>omography (ReFLeCT), which we applied to perform fluorescence imaging of several freely moving zebrafish and fruit fly larvae. Our algorithm reconstructs not only the 3D fluorescence distribution of the sample, but also its 3D optical attenuation map, which is possible due to the extreme angular coverage of ReFLeCT. To account for rapid changes in animal position and orientation throughout the videos, we developed a tracking and registration algorithm that enables fully 3D observation of physiological properties (e.g., morphological dynamics, eye movement, cardiac function) that would otherwise require immobilizing the animal to accurately record. We also demonstrate multi-organism imaging (Fig. 1c, Supplementary Fig. S4, Videos 4 and 5), opening the door to high-speed volumetric imaging of behavioral interactions among multi-mm-sized organisms [28–32].

**Fig. 1.**
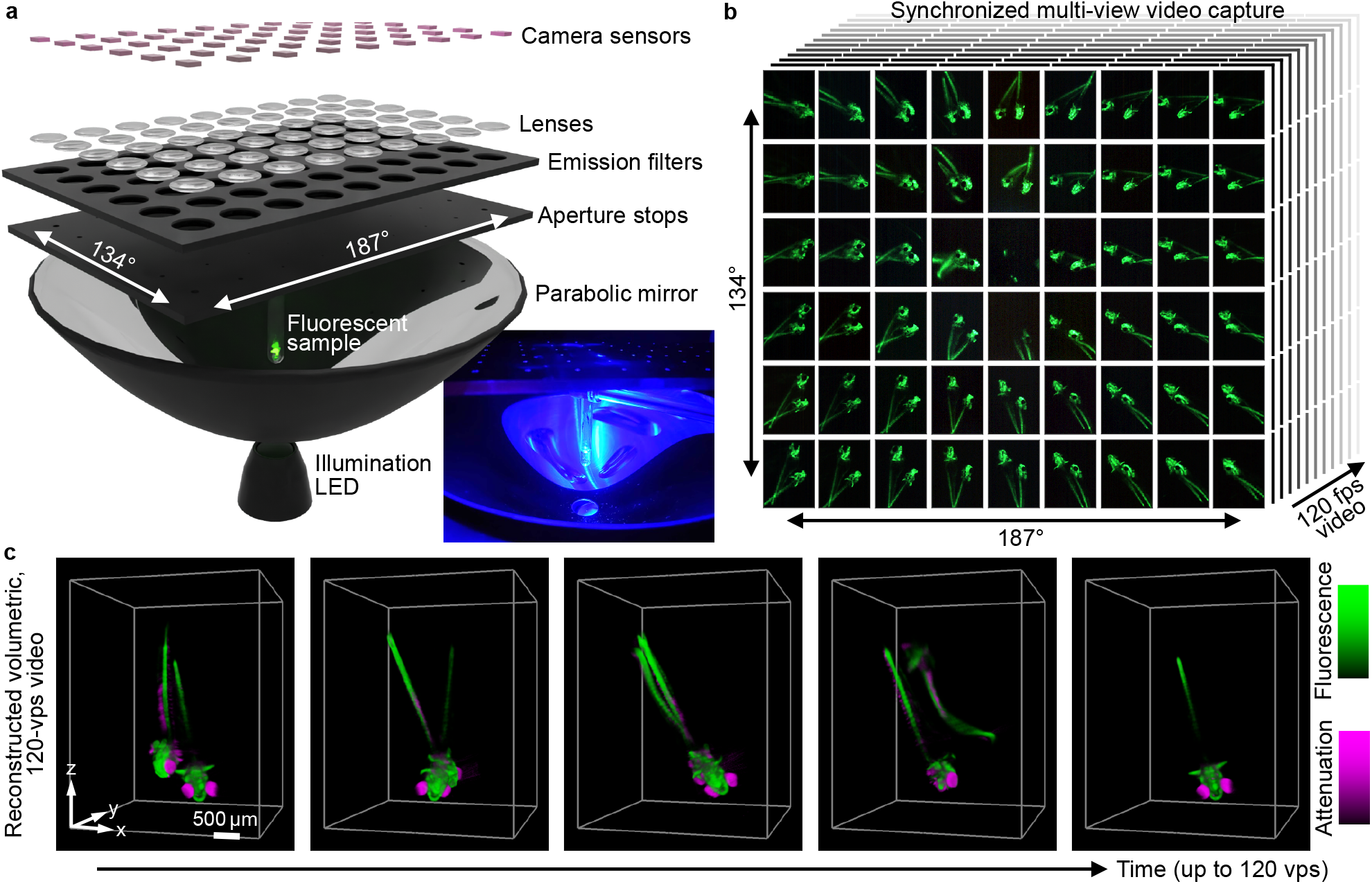
Overview of ReFLeCT. **a** System design, consisting of an array of 54 multi-view fluorescence imaging systems enabled by a large parabolic mirror objective, with specimen placed at the parabola focus. Inset, photograph of a sample under blue LED illumination. **b**, ReFLeCT captures 54 synchronized multi-view videos, spanning an extremely wide angular range (nearly 2*π*steradians), at up to 120 fps, limited by the camera readout rate. **c**, The ReFLeCT computational reconstruction algorithm reconstructs 3D volumetric videos with two channels, fluorescence and opacity (attenuation coefficient), and a volume rate of up to 120 volumes/sec with near-isotropic 3D resolution. See Video 1 for a dynamic overview.

## 2 Reflective Fourier light field computed tomography (ReFLeCT)

### 2.1 Concept and design

ReFLeCT combines a parabolic mirror with a multi-camera array architecture, thereby enabling snapshot multi-view spatioangular sampling of a given volume of interest with significantly higher sampling than conventional Fourier light field microscopes (Fig. 1; Video 1). Working backwards from the plane of image capture, the ReFLeCT system consists of an array of 54 camera sensors (ONSemi AR1335, 3120 *×* 4208 pixels each, 1.1-µm pitch) arranged in a 9 by 6 grid, with a center-to-center camera spacing of *p* = 13.5 mm (Ramona Optics Inc.). We are able to stream data from the entire array (up to 700 megapixels per snapshot, synchronized across all 54 image sensors) to computer memory at rates exceeding 5 GB/sec.

Each camera sensor has an identical lens (Edmund Optics) whose principal planes are approximately a focal length distance *f*_*lens*_ away, so that the object planes are at infinity. Another focal length distance below the lens principal planes (i.e., the Fourier planes) is an array of circular apertures, which serve as the aperture stops that define the lateral resolution and depth of field (DOF) of the system’s acquired multiview images (Sec. 2.2). Between the apertures and lenses is an array of bandpass emission filters (Chroma, 530/50 nm) for green fluorescence imaging. Finally, below these arrays is a large, rotationally-symmetric parabolic mirror (Optiforms, Inc.) with a *f*_*mirror*_ = 25.4-mm focal length and 141-mm-diameter aperture, which acts as a common reflective objective for all 54 cameras. Thus, the 54 camera lenses can be thought of as tube lenses, forming nearly-4f imaging systems with the parabolic mirror acting as the primary objective lens, thereby enabling multi-view imaging spanning *∼*2*π* steradians of a sample placed at the focus of the mirror. The sample is illuminated through a 12.7-mm-diameter hole at the apex of the parabolic mirror with a blue LED (Thorlabs, *λ*=455 nm) with a 500-nm short-pass excitation filter (Thorlabs).

The sample holder is an important design consideration for the imaging system, as it not only holds the sample and impacts the FOV, but also influences the image quality, as it acts as the first optical element of the ReFLeCT system after the sample (see Sec. 2.3). Ideally, the sample holder is a spherical shell with uniform thickness (i.e., an optical dome), which introduces minimal aberrations and allows unobstructed observation from any view angle [25]. To this end, we adopt tubes designed for chemical analysis by nuclear magnetic resonance (NMR) spectroscopy, which have hemispherical bottoms that transition into a cylindrical shaft, and are produced to have as uniform wall thickness as possible to avoid wobble while spinning.

### 2.2 System properties: angular coverage, resolution, magnification, fields of view

The key parameters that determine ReFLeCT’s angular coverage, resolution, magnification, and FOV are *f*_*mirror*_, pinhole aperture diameters, and the lateral positions of the cameras. Due to the simple, transparent geometry of our system, it is straightforward to establish these relationships.

The multiview angular coverage of ReFLeCT is determined by *f*_*mirror*_ and the sensor array spacing and dimensions. In particular, each camera images the sample from a different inclination angle *θ*, dictated by the radial position *r* of the sensor across the mirror aperture, according to the following equation [25]:

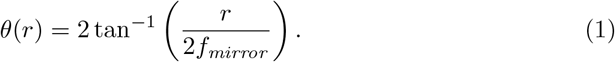

The outer edge sensors in our system had radial*√*positions ranging from *r* = 2.5*p* = 33.75 mm (the shorter array dimension) to 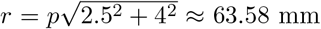 mm (the corner cameras), corresponding to *θ* = 67.12^*°*^ and 102.84^*°*^. In total, our system covers a solid angle of 1.814*π* or nearly 2*π* steradians. For comparison, the solid angular coverage of a 0.90-NA air objective and a 1.10-NA water immersion objective is 1.128*π* and 0.876*π* steradians, respectively.

The lateral resolution of the multiview images is determined by the entry position (*r*) across the mirror aperture and the pinhole diameter. In particular, the object-side effective focal length varies with *r* like [25]

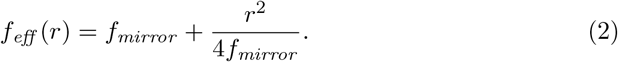

Using this equation, we calculated the pinhole aperture diameters necessary for a view-angle-independent lateral resolution of ~16 µm, which ranged from 0.8 to 2 mm. Another consequence of the positionally-varying effective focal length is that the 54 imaging paths with identical tube lenses can’t simultaneously be 4f systems. Thus, each image has a different magnification, given by

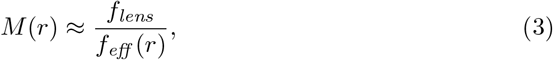

and a different degree of telecentricity, both of which were pre-calibrated prior to sample reconstruction (Methods Sec. 5.1 and Supplementary Sec. S2).

For a given camera, the FOV depends not only on the magnifications and sensor size, but also the lateral resolution or NAs, tuned by the aperture sizes. Assuming the magnification of the image on the sensor is not the limiting factor, the lateral FOV is restricted by the tilt aberrations of the parabolic mirror, which is inversely related to *NA* or *NA*^2^, depending on *r* [25]. Notably, the *NA*^*−*2^ scaling is analogous to the defocus aberration induced by the DOF. In other words, the mirror aberrations do not add additional constraints to the FOV over those enforced by the DOF (see details in Ref. [25]). Thus, the optically-limited volume-bandwidth product, or effective number of resolvable 3D points, of ReFLeCT increases with decreasing NA, creating a unique trade-off space for future design optimization efforts (see Discussion).

### 2.3 Computational optical modeling and reconstruction

We modeled ReFLeCT’s image formation process using a ray-based forward model (Fig. 2), propagating chief rays between the sensors and the sample through all the optical components (similar to Ref. [27]). Although forward models typically start at the sample and terminate at the sensor to predict the measured data, here, we propagated rays in the reverse direction to ensure Cartesian pixel sampling of each video frame. We started propagation of the chief rays corresponding to each pixel of each sensor from the centers of the pupil planes (the stop), where they all intersect, on a per-camera basis. Thereafter, the rays were propagated (Fig. S1) 1) to the parabolic mirror surface and reflected, 2) to the NMR tube surface and refracted, 3) through the NMR tube wall to the glass-water interface and refracted, and 4) through the water in which the biological sample is freely moving.

**Fig. 2.**
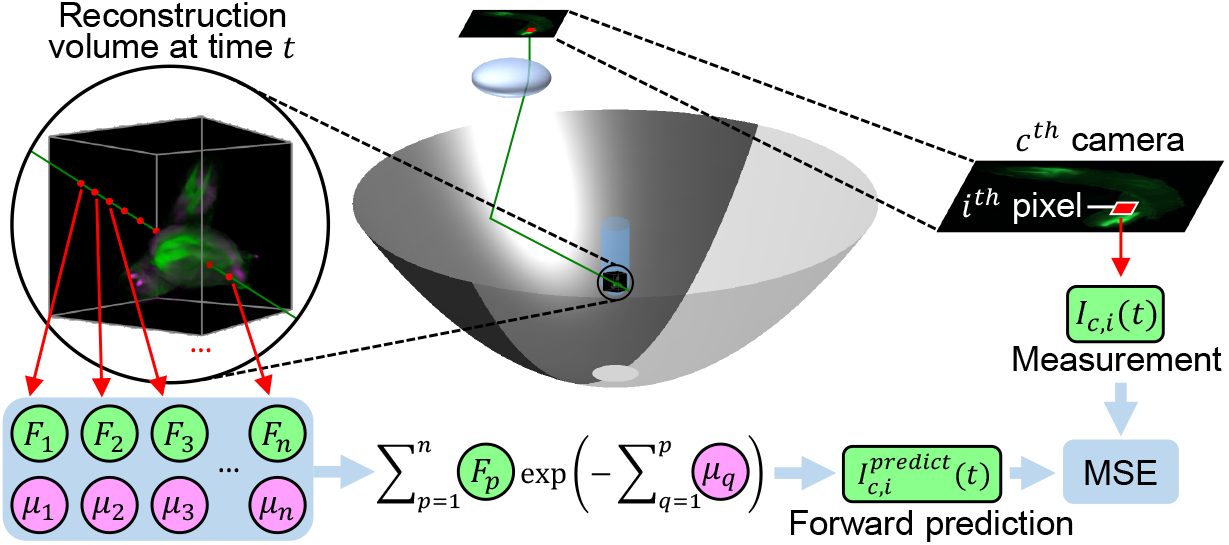
Overview of forward model of the reconstruction algorithm post system calibration. For clarity, a single chief ray from one pixel (*i*) of one camera (*c*) at time *t* is shown, which propagates through the sample, represented as a two-channel 3D volume. For each time point, the reconstruction volume is sampled at *n* coordinates to obtain a list of the fluorescence (*F*) and attenuation (*μ*). These values are passed through the Beer-Lambert law to predict the measured intensity at the pixel, the difference between which are used to compute the mean square error (MSE), which in turn is minimized via stochastic gradient descent (SGD) with respect to the reconstruction. This reconstruction procedure is repeated for all time points.

In practice, we also accounted for known sources of misalignment parametrically via translation vectors and rotation matrices, such as the relative pose between the camera array and parabolic mirror and the pose of the NMR tube. While such parametric modeling promoted convergence by offering a good initial guess, alone it was insufficient to accurately predict our data, so we enriched our ray propagation model nonparametrically using high-order polynomials to account for unknown sources of misalignment and distortions (Supplementary Sec. S1.2). For a detailed mathematical treatment of the full ray propagation trajectory, see Supplementary Sec. S1. These calibration parameters are estimated in a preceding, multi-step calibration procedure (Methods Sec. 5.1) prior to reconstruction.

Once we obtain the final calibrated rays, specified by position vectors 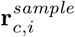 and direction unit vectors 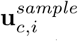, for every pixel *i* across every camera *c*, we sampled the dynamic 3D fluorescent object *F* (**r**, *t*), along the ray trajectories (Fig. 2). To account for attenuation caused by absorption or scattering, we simultaneously modeled a coregistered dynamic 3D attenuation distribution *µ*(**r**, *t*) of the object and also sample voxels along the ray trajectories. To obtain the forward prediction of the pixel *i* of camera *c*, we summed the sampled fluorescence values along the ray, modulated by the attenuation values, in accordance with the Beer-Lambert law (Methods Sec. 5.2). Finally, we iteratively reconstructed *F* (**r**, *t*) and *µ*(**r**, *t*) by minimizing the mean square error (MSE) between the forward prediction and the measured pixel intensity using stochastic gradient descent (SGD) for each time point independently (Methods Sec. 5.3).

## 3 Results

### 3.1 Resolution and FOV characterization

To characterize the resolution and FOV of our ReFLeCT system, we imaged a sparse distribution of 6-µm green fluorescent beads, embedded in 1% agarose in an NMR tube. We reconstructed the fluorescent beads on an isotropic voxel grid with a voxel size of 8 µm (Fig. S3a,b). We then segmented the beads and performed fitting using a separable 3D Gaussian model, and reported the full widths at half maximum (FWHMs) (Fig. S3c,d). As expected, the resolution is best at the bottom of the tube (15-20 µm), where the walls form a spherical shell and thus introduce minimal aberration. The resolution gets worse the further away from the bottom of the tube, due to astigmatism from the cylindrical walls and increased distance from the nominal focus of the parabolic mirror, which leads to more defocus and intrinsic tilt aberrations from the parabolic mirror [25]. Nevertheless, the resolution is nearly isotropic, regardless of the position within the FOV (Fig. S3d). The 3D FOV of this implementation of ReFLeCT is thus limited by the minimum acceptable resolution. If we assume a resolution cutoff of *∼*30 µm, our effective FOV volume would cover *∼*25 mm^3^, while a more generous cutoff would yield *∼*41 mm^3^.

### 3.2 Fruit fly larvae (*Drosophila melanogaster*)

Next, we applied ReFLeCT to image several freely moving transgenic fruit flies at late larval stages (3rd instar wandering larva, WL3), expressing GFP in their muscle cells (*sqh-GFP*, Fig. 3; Video 2), pericardial cells (*HandC-GFP*, Fig. 4; Video 3), and salivary glands (*NP5169 Gal4>UAS-GFP-NLS*, Fig. S4; Video 4). The volumetric frames of these videos were reconstructed with a voxel size of 16 µm, twice as large as that for the beads (Sec. 3.1) to boost SNR. Thus, our 3D spatial resolutions are voxel-limited.

**Fig. 3.**
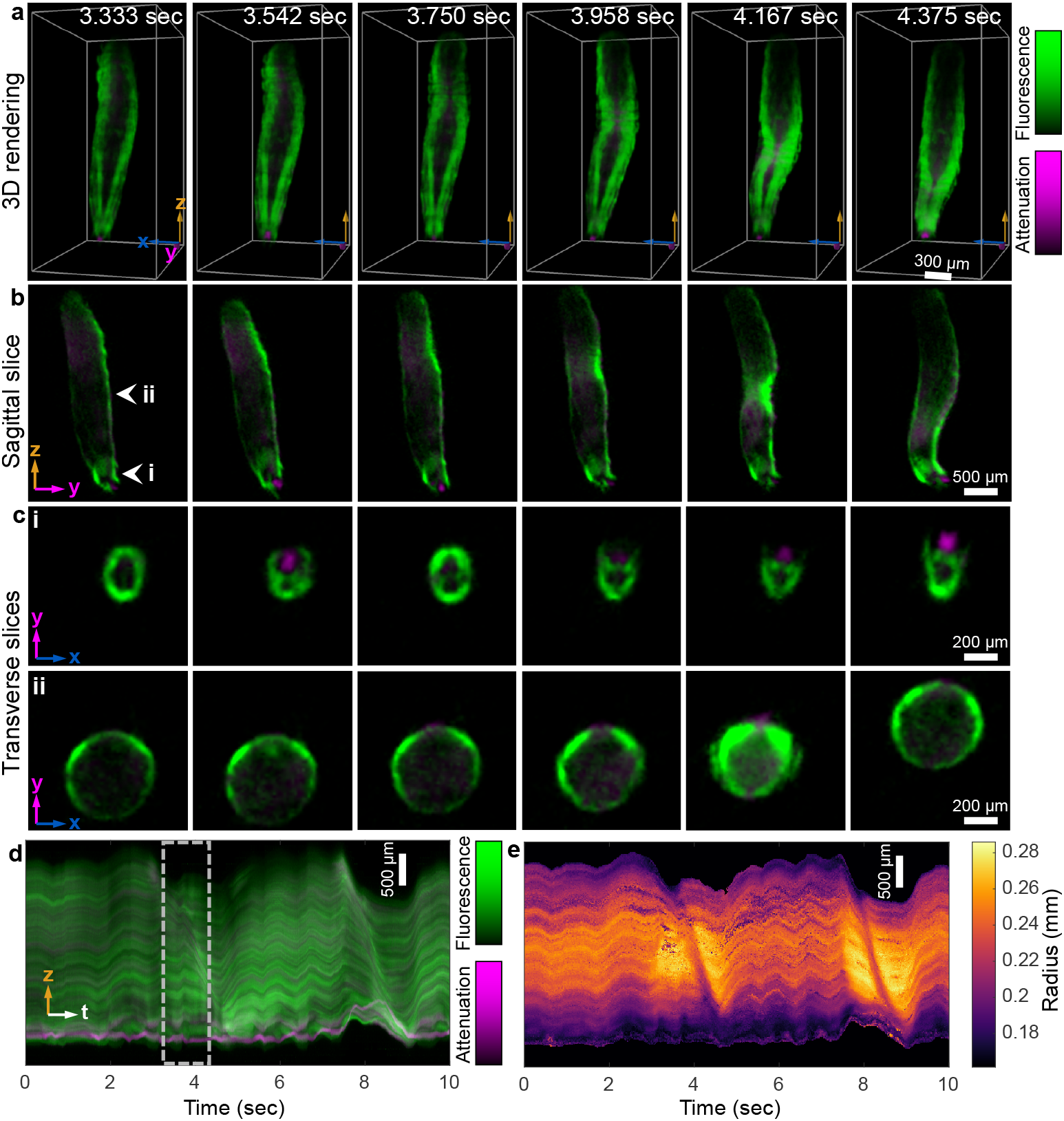
Volumetric imaging of a freely moving WL3 *Drosophila* larva (*sqh-GFP*) with GFP-labeled myosin at 120 vps. See also the associated Video 2. **a**, 3D renderings from a fixed perspective of the larva at several time points throughout a segmental muscle contraction. **b**, Sagittal (*yz*) cross-section of the larva over time. **c**, Two transverse (*xz*) cross-sections (**i** & **ii**, indicated by white arrowheads in **b**) over time. The first row shows the jaw motion. **d**, Summary kymograph of the 10-sec video: 1D max intensity spatial projections across *x* and *y*, plotted across time. Dotted box indicates the time range during which the segmental muscle contraction illustrated in **a**-**c** occurs. Another contraction occurs around 8 sec. **e** Radius of the larva across the length of its body and time.

**Fig. 4.**
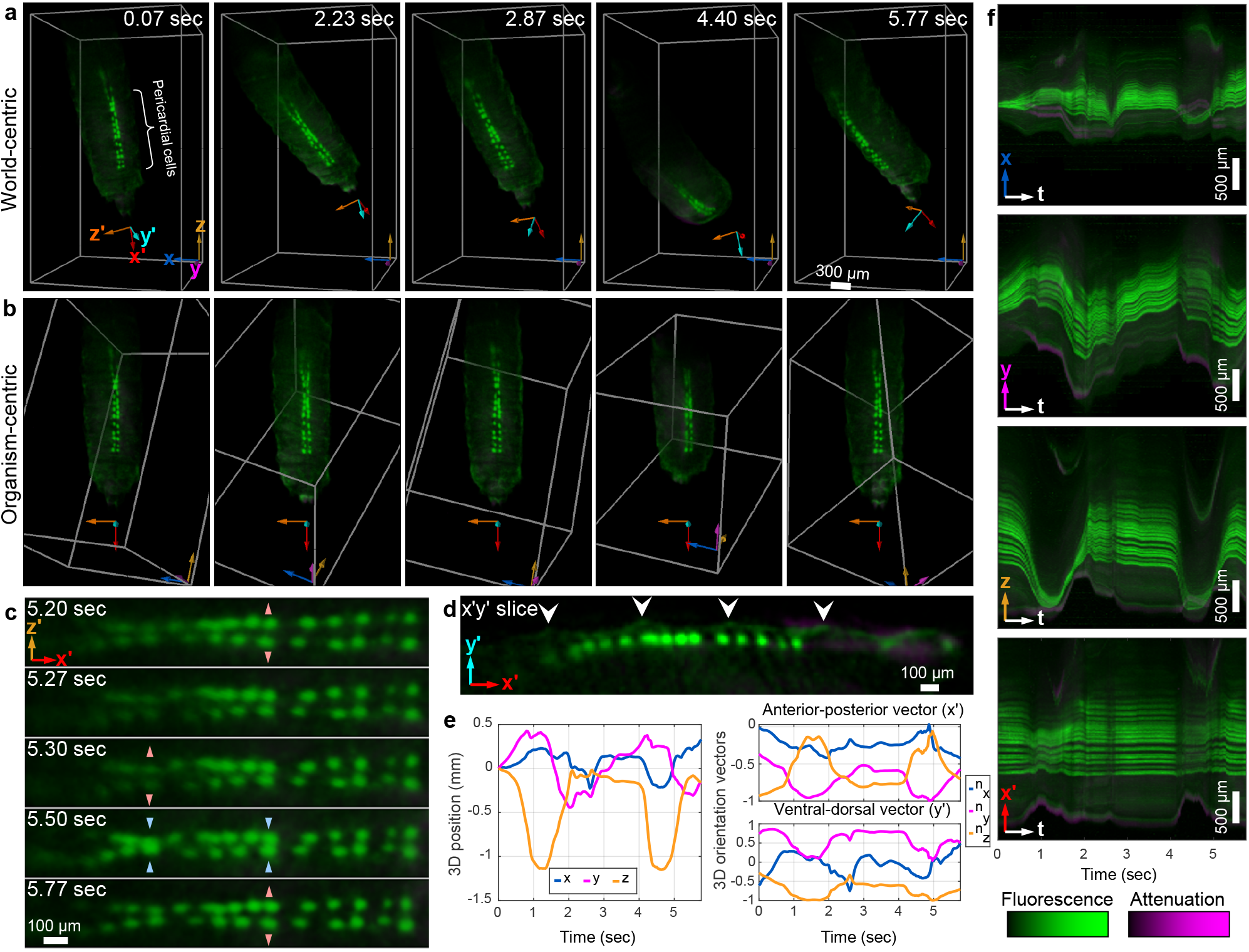
Volumetric imaging of a freely moving WL3 fruit fly larva (*HandC-GFP*) with GFP-labeled pericardial cells at 30 vps. See also the associated Video 3. **a**, 3D renderings from a fixed perspective of the larva at several time points. The primed coordinate system is relative to the larva, while the unprimed coordinate system is of the imaging system (“world”). **b**, 3D renderings at the same time points as in **a**, but with a perspective locked onto the larva. **c**, Close-up of the pericardial cells at five time points. Arrowheads indicate when a pair of pericardial cells are moving closer to each other or further away. **d**, Sagittal cross-section through the larva, showing that the pericardial cells are *∼*50 *μ*m below the surface. White arrowheads indicate the segmental furrows interleaving the body segments. **e**, 6D pose of the larva over time. Left plot is of its 3D position, referenced to its initial position. Right plots show 3D orientation, represented by two larva-centric unit vector axes (*x′* and *y′*, shown in **a**). **f**, Visualization of the 4D data (3D + time), max-intensity-projected down to 2D kymographs (*xt, yt*, and *zt*). The fourth kymograph (*x′t*) projects across the *larva*’s axes (*y′* and *z′*), with residual motion due to the larva changing its curvature.

From the reconstructed 10-sec, 120-vps video of the *Drosophila* larva with GFP-labeled muscle (*sqh-GFP*), summarized in Fig. 3 and Video 2, we observed two peristaltic muscle contraction events, manifesting as a local constriction propagating from the posterior to anterior of the larva (Fig. 3a), lasting about 1 sec in duration [33–35]. This localized constriction results in increased localized fluorescence density (Fig. 3b,c(ii)). During the contractions, the length of the larva also decreases before it increases upon uniform length-wise relaxation (see kymograph in Fig. 3d, 3.5-4.5 sec and 7.5-9 sec). ReFLeCT also enabled us to track the radius of the larva’s transverse cross-sections along the length of the body and over time (Methods Sec. 5.8), confirming the radius decrease at the point of constriction as it propagates during the segmental muscle contraction (Fig. 3e). Finally, we also observed jaw motion throughout the contraction (Fig. 3b,c(i); Video 2, sagittal and transverse slices).

Next, we reconstructed a 30-vps volumetric video of a freely moving *Drosophila* larva with GFP-labeled pericardial cells (*HandC-GFP*, Fig. 4; Video 3). Since the larva significantly changed its 6D pose (3D position and 3D orientation) throughout the video (Fig. 4a), we tracked the larva (Methods Sec. 5.7) and digitally repositioned and reoriented the virtual camera view so that larva was always in the same pose, with the pericardial cells facing the virtual camera (Fig. 4b). We have thus defined two coordinate systems: the “world” coordinate system, where *z* points towards the camera array, and the organism coordinate systems, denoted with primes, where *x*′ points from anterior to posterior, *y*′ points from ventral to dorsal, *z*′ from right to left. As a result, the world coordinate system remains static in Fig. 4a and reorients across time in Fig. 4b, the opposite is true for the organism-centric coordinates. The 6D tracking results are summarized in Fig. 4e, which shows the 3D position of the larva relative to its initial position, and the directions of the *x*′ and *y*′ unit vectors. The dynamic nature of the larva and its subsequent tracking can also be visualized in the kymographs in Fig. 4f, where the kymograph post tracking is more stabilized, with some residual deformations of the longitudinal pericardial cell arrangement, due to the larva changing its curvature.

From the tracked larva, we were able to observe dynamic changes in the spacing of the pericardial cells over time (Fig. 4c), due to its proximity to the heart [36]. The tomographic imaging capabilities of ReFLeCT also enabled us to observe that the pericardial cells are *∼*50 µm below the surface of the larva (Fig. 4d).

Finally, we demonstrated multi-organism imaging by reconstructing a 10-sec, 30-vps video of two fruit fly larvae with GFP-labeled salivary glands (*NP5169 Gal4>UAS-GFP-NLS*, Fig. S4; Video 4). The tomographic imaging capabilities of ReFLeCT also recovers the hollow tube shape of the salivary glands (Fig. S4d) [37].

### 3.3 Zebrafish larvae (*Danio rerio*)

We also applied ReFLeCT to image several freely swimming zebrafish larvae, including one at 7 days post fertilization (dpf) expressing GFP in plasma membranes in the jaw and notochord (*nfatc:gal4;UAS:GFP-CaaX*) at 120 vps (Fig. 5; Video 5), one at 4 dpf expressing GFP in the heart (*cmlc2:GFP*) at 30 vps (Fig. 6; Video 7), and another at 6 dpf expressing GFP in neuronal plasma membranes (*gad1b:gal4;UAS:GFP-CaaX*) at 120 vps (Video 6). The reconstruction voxel size (16 µm) and larva-centric coordinate systems are the same as the ones we used for fruit fly larvae (Sec. 3.2).

**Fig. 5.**
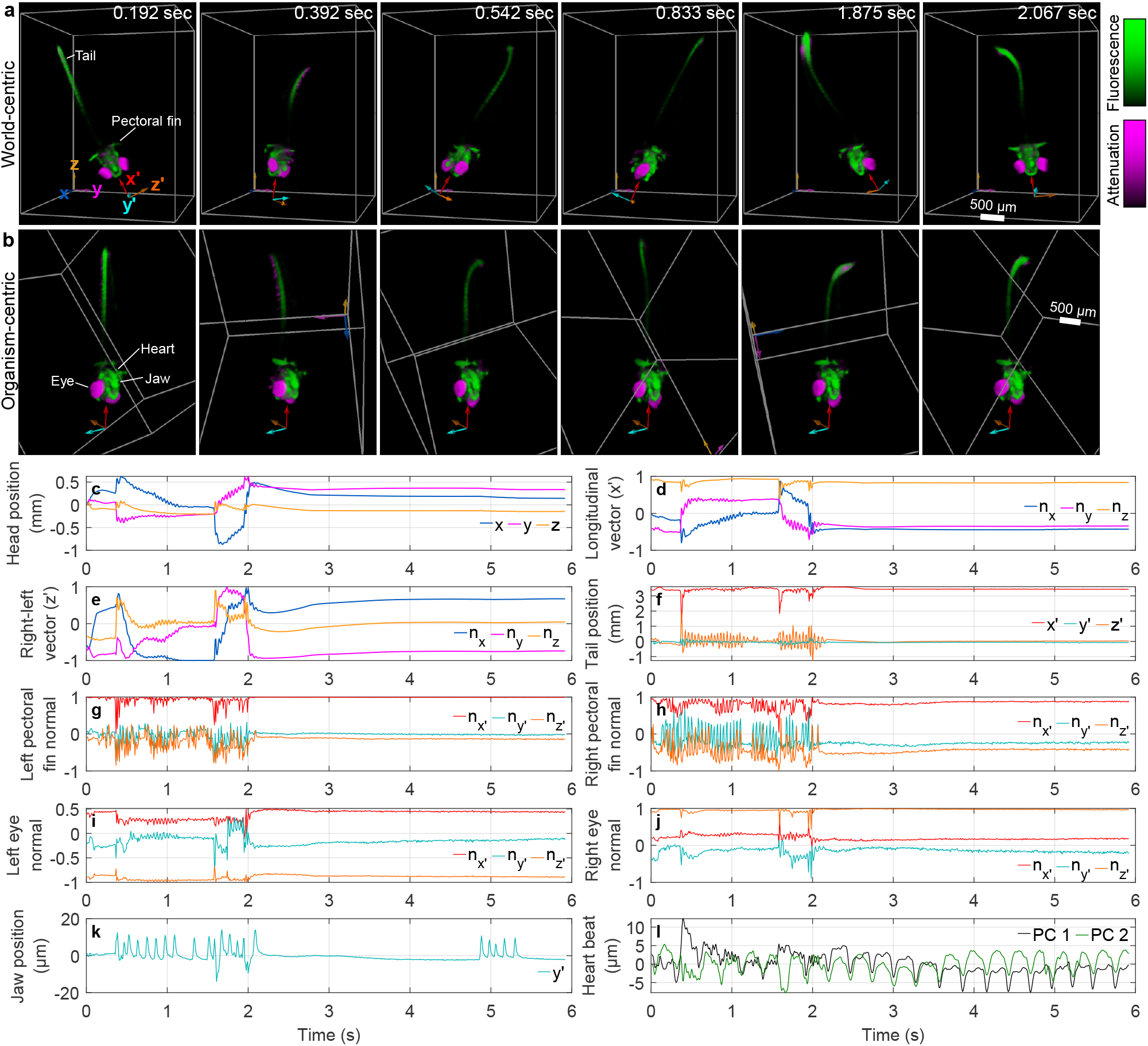
Volumetric imaging of a freely moving 7-dpf zebrafish larva (*nfatc:gal4;UAS:GFP-CaaX*) at 120 vps, with GFP expression in plasma membranes in the jaw and notochord. See also the associated Video 5. **a**, 3D renderings of the larva from a fixed perspective over time. The primed axes are relative to the larva, while the unprimed axes are of the imaging system (“world”). **b**, 3D renderings, perspective-locked to the larva’s tracked head. **c**-**k**, Various tracked properties of the fish over time. The color-coding matches that of the axes in **a**, depending on whether they reference the larva’s or the world coordinate system. **c**, 3D position of the larva’s head over time, relative to its initial position. **d**-**e**, The 3D orientation of the larva over time, represented by the components of the unit vectors pointing from its anterior to posterior (*x′*) and its right to left (*z′*). **f**, Tail position of the larva over time, relative to the tracked head position. **g**-**h**, Orientation of the left and right pectoral fins, respectively, represented as the unit vector normal to the fin. **i**-**j**, Gaze direction of the left and right eye, respectively. **k**, Jaw position over time. **l**, Tracked heart beat over time, represented as the first two principal components (PCs) of its motion.

**Fig. 6.**
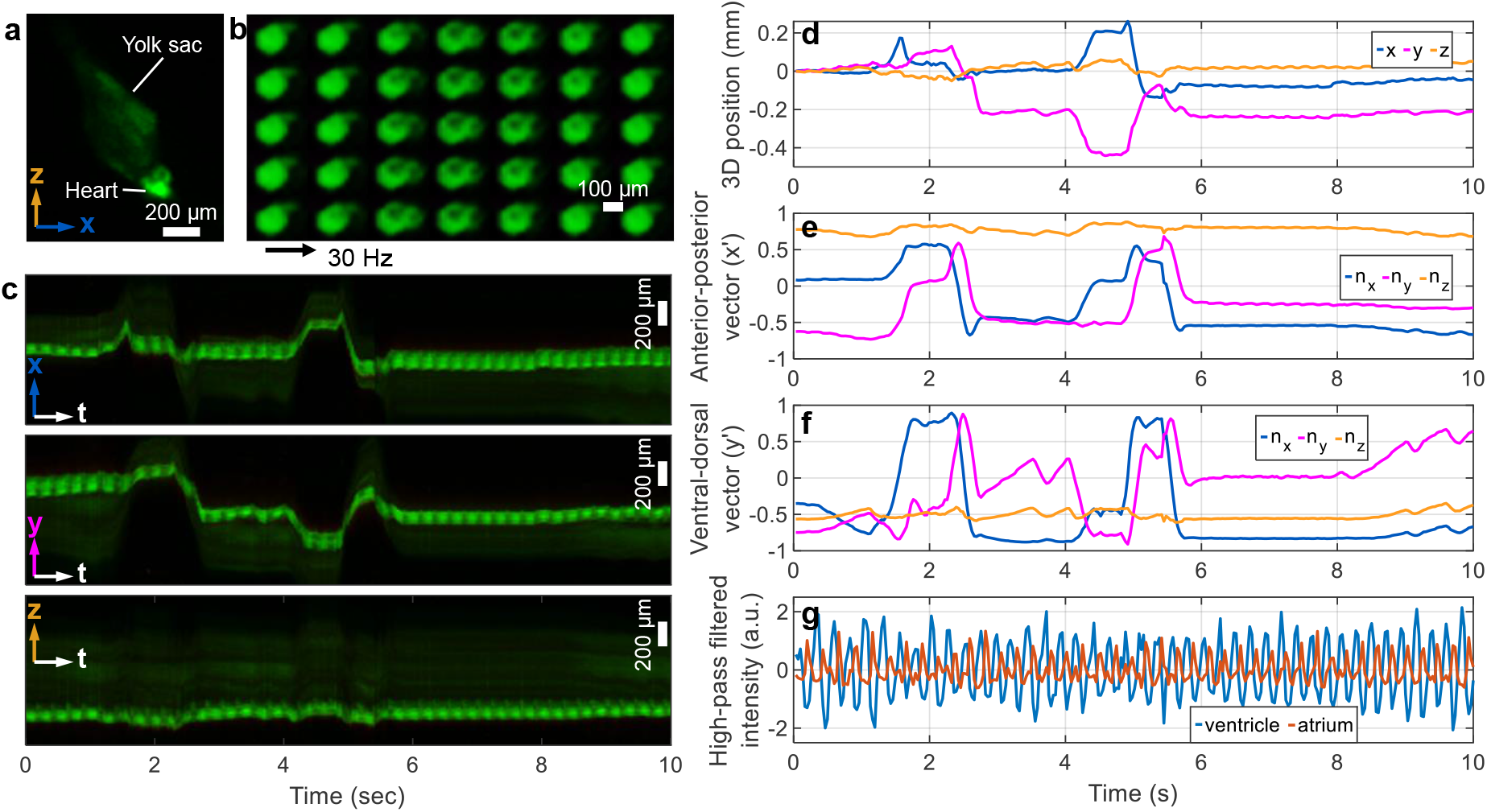
Volumetric imaging of a 4-dpf zebrafish larva (*cmlc2:GFP*) expressing GFP in the heart, at 30 vps. See also the associated Video 7. **a** *xz* max projection of 3D reconstruction at one time point. **b**, Six sequential cardiac cycles, showing its two heart chambers (ventricle and atrium). The time difference between sequential images is 33 ms. **c**, 1D max projections across two of the three spatial dimensions, plotted across time. The *x*/*y*/*z* axes are relative to the imaging system (i.e., not the fish). **d**, The 3D position of the fish over time, relative to its position at the start of the video. **e**-**f**, The 3D orientation of the fish over time, represented by the unit vectors pointing from head to tail (**e**) and from its ventral to dorsal sides (**f**). **g**, Temporally high-pass filtered fluorescence intensity of the ventricle and atrium.

Fig. 5a shows a few reconstructed volumes of the 120-vps video of the 7-dpf zebrafish larva (*nfatc:gal4;UAS:GFP-CaaX*). Here, not only does the green fluorescence provide useful contrast, but also the attenuation channel (magenta) highlights the eyes of the larva as well as the pigmentation patterns spanning the length of the body. Since the zebrafish larva rapidly changed its position and orientation as it swam, we performed 6D pose tracking of the head, using only the green fluorescence channel (Methods Sec. 5.7). The tracking data is summarized in Fig. 5c-e, showing the 3D head position, and the larva’s ventral-dorsal (*x*′) and right-left (*z*′) axis orientation plotted against time. Using this tracking data, like with *Drosophila* larva (Fig. 4, Sec. 3.2), we dynamically changed the virtual camera perspective throughout the reconstructed video so that the fish head appears static (Fig. 5b).

Since we tracked the 6D pose of and coregistered the larva from all time points, we were able to track other many other properties of the larva (Fig. 5f-l). One property was the 3D position of the tail tip (Fig. 5f), which moves primarily in the *z*′ axis (left-right), and occasionally produces large strokes that result in the tail getting close to the head (near 0.4 and 1.6 sec). We also tracked the orientation of the left and right pectoral fins (Fig. 5g,h), as well as the gaze of both eyes based on the attenuation channel (Fig. 5i,j). Note that the gaze oscillates along with the rest of the body, possibly due to the vestibulo-ocular reflex (VOR) [38, 39]. These oscillations are not an artifact of the volume registration, as only the fluorescence channel was used for registration. We also tracked the jaw motion (Fig. 5k) and the heart beat of the fish (Fig. 5l), which are both visible in Fig. 5b (and Video 5).

Additionally, we imaged a 4-dpf zebrafish larva at 30 vps, with GFP labeling localized exclusively to the heart (*cmlc2:GFP*, Fig. 6a; Video 7). The kymographs in Fig. 6c show the trajectories of the freely swimming larva. Upon 6D pose tracking (Fig. 6d-f) and volumetric coregistration, we were able to visualize the atrium and ventricle of the heart (Fig. 6b), including the phase offset delay between their contractions (Fig. 6g).

## 4 Discussion

Reflective Fourier light field computed tomography (ReFLeCT) is a new high-throughput computational imaging technique for tomographic 3D video over 10s-of-cubic-mm volumes at up to 15-20-µm isotropic 3D spatial resolution and 120-Hz synchronized frame rates, culminating in multi-gigavoxel/sec dynamic volume throughputs. Using the high speeds and large viewing volumes of ReFLeCT, we demonstrated behavioral imaging of freely moving organisms (*Drosophila* and zebrafish larvae) without the need for anesthesia or restraint, thus avoiding confounding effects on organism behavior and physiology [19, 40, 41]. To elucidate and quantify their unconstrained 3D behavior, we developed a 6D pose tracking and registration algorithm for perspective-locking onto the animal. With this software, we were able to track and quantify physiological dynamics such as heart beat, eye gaze, and fin motion, all in full 3D, which otherwise require organism immobilization for longitudinal measurement. ReFLeCT also opens up the possibility of high-speed volumetric video observation of behavioral interactions across multiple organisms - a new capability with relevance in neuroscience and developmental biology, for example [28–32].

We believe ReFLeCT is a significant step forward for high-speed volumetric video microscopy, with several avenues for improvement. While the resolution of our prototype is isotropic and sufficient for behavioral imaging, higher resolution may be necessary for a variety of functional imaging applications, in particular to monitor neural activity. Since ReFLeCT’s resolution is tied to the DOF of each imaging module and therefore its total volume of view, it would be straightforward to design new ReFLeCT systems that capture across smaller volumes at higher resolution (and conversely, larger volumes at lower resolution). Alternatively, one can employ DOF extension strategies [23, 42–45] to achieve higher resolution while maintaining a large volume of view. Increasing the resolution by expanding the aperture sizes of each imaging module would also increase overall measurement SNR by allowing greater light collection efficiency. It is also direct to envision multi-wavelength excitation and emission filtering strategies for multi-channel fluorescence video sampling with ReFLeCT, which could increase cellular-level specificity and allow ratiometric analyses.

While the spatiotemporal throughput of our current ReFLeCT design is limited by the data transfer rate from our sensors to the computer to *∼*5 GB/sec [46, 47], improvements to our data transfer architecture and the creation of image sensor arrays with faster camera sensors could enable applications in imaging fluorescent voltage sensors to monitor neural activity [48] and capture high-speed behaviors (e.g., rapid zebrafish locomotion during swim bouts [49] or kinematics during seizure activity [50]), potentially simultaneously across multiple organisms. On the computational backend, to handle such increased spatiotemporal data throughputs, we could adapt neural radiance fields (NeRFs) [51] and recent advances in accelerating optimization speed [52] and extending them to implicit 4D representations [53–57]. Moreover, reparameterizing tomographic reconstructions as outputs of neural networks can offer regularizing effects [58, 59]. With such improvements, ReFLeCT could readily record high-speed 3D fluorescent dynamics within a variety of alternative organisms (e.g., *C. elegans* [60], jellyfish [61], 3D cellular models (e.g., neural activity within cerebral organoids [62]), and alternative setups (e.g., 3D flow cytometry [63]) to open up new avenues for future scientific exploration.

## 5 Methods

### 5.1 Imaging system calibration

To account for optical misalignment, manufacturing imperfections, and distortions in our ReFLeCT system, we performed a three-step calibration procedure prior to sample reconstruction:

1. Calibration of the camera array and mirror without the NMR tube, using precise translation of a fiber optic cannula with a diffuser tip and a backwards bundle adjustment algorithm. See Supplementary Sec. S2 for details.
2. Coarse estimation of the NMR tube pose based on a freely-moving organism’s exploration of the tube lumen. See Supplementary Sec. S3 for details.
3. Joint refinement of sample-incident rays and estimation of a low-resolution reconstruction of *F* (**r**, *t*) for multiple values of *t*. See Methods Secs. 5.2 and 5.3 about the tomographic reconstruction procedure for more details.

While most prior works require immobilization of the organism, here, in the second and third steps, we actually rely on the organism exploring broadly within the NMR tube, which improves the posedness of the tube calibration.

### 5.2 Forward modeling

Following calibration of the ray trajectories at the sample for each pixel *i* across each camera *c*, parameterized by position 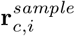 and direction 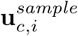, we summed the resulting fluorescence values (*F* (**r**, *t*)) along the rays, subject to modulation by the attenuation values (*µ*(**r**, *t*)) along the rays, to obtain the forward prediction,

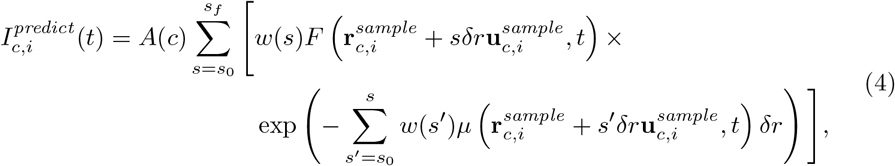

following the Beer-Lambert law, where *δr* is the discretization step size for ray propagation and *s* is the number of discrete steps, ranging from *s*_0_ to *s*_*f*_, *A*(*c*) is a per-camera scale factor to account for amplitude variation (e.g., due to camera-dependent magnification), and *w*(*s*) is a window function that enforces a soft constraint on the summation range,

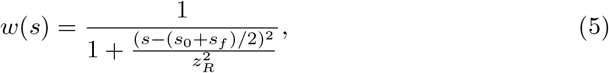

modeling the peak intensity of a Gaussian beam along the propagation direction, with DOF *z*_*R*_.

As *F* (**r**, *t*) and *µ*(**r**, *t*) were modeled discretely via a voxel grid for each time point, the input spatial coordinates were interpolated into the nearest 2*×* 2*×* 2 voxels to allow gradients to propagate through all the calibration parameters for gradient descent optimization. This interpolation is only necessary during calibration (i.e., step 3 of Methods Sec. 5.1) and not during the final object reconstruction, for which we simply rounded to the nearest voxel.

### 5.3 Inverse 4D reconstruction

To reconstruct dynamic tomograms of the 3D objects within the sample holder, we developed a reconstruction algorithm that computes a weighted mean square error (MSE) between the forward prediction of camera pixel intensities (Eq. 4) and the actual measured data,

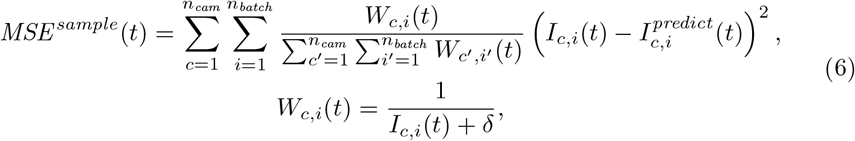

where *n*_*cam*_ *≤*54 is the number of cameras and *n*_*batch*_ is the number of pixels randomly selected across each camera sensor per batch during mini-batch stochastic gradient descent (SGD). Each batch consists of propagating a total of *n*_*cam*_ *n*_*batch*_ rays for a given time point (*t*). The weight function *W*_*c,i*_(*t*) gives greater emphasis to dark background pixels to reduce reconstruction artifacts outside of the organism. When *δ* is close to 0, dark background pixels are weighted more heavily, while when *δ* is large, all pixels are weighted approximately equally. In practice, we also precompute a low-resolution occupancy grid, indicating whether each voxel belongs to (or is close to) the organism being imaged, to accelerate convergence (Supplementary Sec. S4).

We did not adopt spatial smoothness-enforcing regularization methods on *F* (**r**, *t*) or *µ*(**r**, *t*), such as total variation (TV), to avoid blurring and spatial resolution loss. Rather, we only used 3D support constraints – a hard constraint using the occupancy grid (Supplementary Sec. S4) on both *F* (**r**, *t*) and *µ*(**r**, *t*), and an additional soft support constraint on *µ*(**r**, *t*), by including a new weighted L1 regularization term based on points sampled along ray trajectories for the current batch,

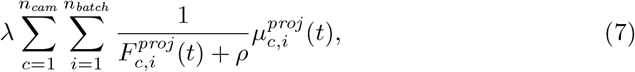

where

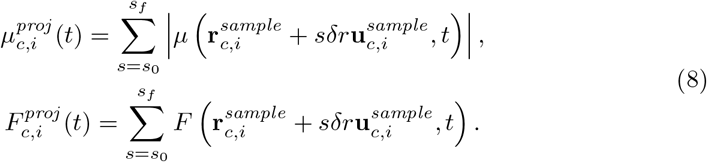

Here, the weighting scheme is similar to that of Eq. 6, where the L1 loss on *µ*(**r**, *t*) along ray trajectories is weighted by the inverse of the projections of *F* (**r**, *t*) along the same ray trajectories, where *ρ* plays a similar role to *δ* in Eq. 6 and *λ* tunes the strength of the regularization. Thus, this regularization term applies shrinkage to *µ*(**r**, *t*) more strongly in regions outside of the moving organism. In this way, both Eqs. 6 and 7 offer object support regularization. The reason why we don’t use *I*_*c,i*_(*t*) in Eq. 7 as we did in Eq. 6 is that dark regions in the raw data can be due to either no sample, or a heavily attenuating sample. By using 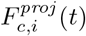, we eliminate the possibility of the latter. Lastly, since 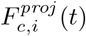 serves as a weight factor, we do not allow gradients to propagate through to *F* (**r**, *t*) during gradient descent, so that Eq. 7 only regularizes *µ*(**r**, *t*).

We sequentially reconstructed *F* (**r**, *t*) and *µ*(**r**, *t*) for each time point to generate full dynamic volumetric videos (300-1200 frames). Using a Nvidia RTX A6000 GPU, we were able to reconstruct each volumetric frame in 5-15 mins. Finally, we visualized the 4D reconstructions from different perspectives using PyVista [64].

### 5.4 Fruit fly larvae preparation

Fruit flies (*Drosophila melanogaster*) were raised at 25°C on standard fly food provided by Archon Scientific. Fly stocks used in this study are *NP5169 Gal4>UAS-GFP-NLS* [65], *HandC-GFP* [66], and *sqh-GFP* (BDSC 57144)[67]. For imaging, wandering L3 (WL3) larvae were used. To remove the autofluorescence from the food inside the gut, *Drosophila* larvae were starved on agar grape juice plates for 12-24h before imaging.

### 5.5 Zebrafish larvae preparation

Zebrafish (*Danio rerio*) stocks were used and maintained at 28°C and were bred as previously described [68] and in accordance with Duke University Institutional Animal Care and Use Committee (IACUC) guidelines. All zebrafish were raised in a circulating aquarium. Housing tanks held 1–10 fish/L. Male and female breeders from 3 months to 1 year of age were used to generate larvae for all experiments. 4-7 dpf zebrafish larvae were used for this study. The strains used in this study were: *Tg(UAS:GFP-CaaX)*^*pd1025*^, *cmlc2:GFP* (from *Tg(UAS:mcherry)*^*pd1112*^), *TgBAC(nfatc1:Gal4)*^*mu286*^ [69], and *Tg(gad1b/GAD67:Gal4-VP16)*^*mpn155*^ [70].

### 5.6 Imaging

The organisms were immersed in water and transferred into the NMR tubes (Wilmad-LabGlass 524-PP-7-5; ID=3.43±0.19 mm, OD=4.9635±0.0071 mm) via Pasteur pipettes. The NMR tubes were then positioned at the parabolic mirror’s focus and imaged by the ReFLeCT system at 30 fps (30-ms exposure) or 120 fps (7-ms exposure). After the imaging session, we captured a calibration dataset by replacing the NMR tube with a fiber-optic point source and translating it in a 3D grid (Supplementary Sec. S2).

### 5.7 Organism 6D pose tracking and registration

We generated organism-centric videos, whereby the virtual camera view locks onto a part of the organism (e.g., the head) so that it appears to remain stationary throughout the video. Creating these videos requires tracking the 6D pose (3D translation + 3D orientation) of the organism over time. To do this, we first performed rigid volumetric registration between adjacent time points using MATLAB’s `imregtform` function using a pixel intensity-based mean square error metric. In other words, we estimate *T*_*i*+1*→i*_, which is the rigid transformation matrix that registers frame *i* + 1 to frame *i*. We then sequentially registered all the frames of the video to the first frame, using a recursively defined initial guess, obtained by composing the optimized transform of the previous frame with the transform that registers the current frame to the previous frame. That is, if *T*_*i→*1_ registers frame *i* to frame 1, then the initial guess for *T*_*i*+1*→*1_ would be *T*_*i*+1*→i*_*T*_*i→*1_. We can then obtain the dynamic 6D poses of the organisms by specifying an initial 3D coordinate and 3D orientation (e.g., 3 orthonormal vectors) and applying these optimized rigid transformation matrices, *T*_*i→*1_.

### 5.8 Fruit fly larva radius estimation

To compute the radii of the fruit fly larva during muscle contractions (Fig. 3e), we started by estimating the lateral center of mass of the larva’s green fluorescence as a function of axial position. From this curve, we estimated the local orientation of the larva as the vector pointing between the centers of mass of two adjacent axial positions. Using the center of mass and this vector, we virtually sliced the larva and used a circular Hough transform to find the radius of the larval cross-section. We repeated this for all axial positions across all time points.

## Data availability

Select multi-view video data will be made available at TBD.

## Code availability

Python code for 4D reconstruction and interactive visualization will be made available on Github.

## Acknowledgements

Research reported in this publication was supported by the Office of Research Infrastructure Programs (ORIP), Office Of The Director, National Institutes Of Health of the National Institutes Of Health and the National Institute Of Environmental Health Sciences (NIEHS) of the National Institutes of Health under Award Number R44OD024879, the National Cancer Institute (NCI) of the National Institutes of Health under Award Number R44CA250877, the National Institute of Biomedical Imaging and Bioengineering (NIBIB) of the National Institutes of Health under Award Number R43EB030979, the National Institute of General Medical Sciences of the National Institutes of Health under Award Numbers R01GM150666, R01GM118447 and R01GM140138, the National Science Foundation under Award Number 2036439, a Duke-Coulter Translational Partnership Grant, the American Heart Association under Award Number 23POST1013432. This work was also supported in part by the Schmidt Science Fellows, in partnership with the Rhodes Trust.

## Contributions

Conceptualization: KCZ, RH; Methodology: KCZ, RH; Software: KCZ; Formal Analysis: KCZ; Investigation: KCZ, CC, AC, KCL, XY, SX, JB; Data Curation: KCZ; Writing: KCZ, CC, KCL, SX, RH; Visualization: KCZ, JJ, RB, MH; Supervision: DF, MB, RH; Funding Acquisition: DF, MB, RH.

## Supplementary information

### S1 Detailed mathematical treatment of ray propagation from the sensor pixels to the sample

#### S1.1 Ray tracing from the sensor pixels to the parabolic mirror focus

We start at the center point of each pinhole aperture stop, whose position is **R**_*c*_ = (*X*_*c*_, *Y*_*c*_, *Z*), where all chief rays originating from the sensor pixels converge and then diverge, by definition. For now, we assume that *Z* is independent of *c* and that (*X*_*c*_, *Y*_*c*_) form a cartesian grid spaced by *p*. These coordinates are defined such that the origin is at the parabolic mirror’s focus. Let 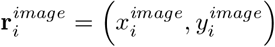 be the nominal relative lateral spatial coordinates of the *i*^*th*^ pixel (assuming identical sensors), defined so that the central camera pixel is (0, 0) and assumed to be a cartesian grid (spaced by the pixel size, 1.1 µm). Then, the nominal ray originating from the *i*^*th*^ pixel at the pupil plane of the *c*^*th*^ camera has position and direction,

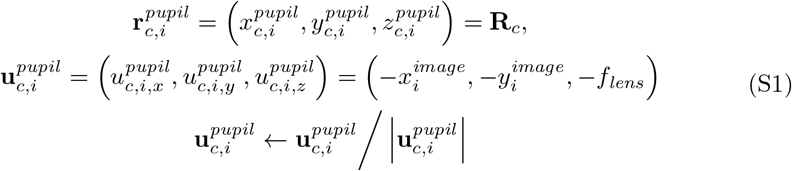

where the +*z* axis points away from the parabolic mirror. We start the ray tracing from this plane and follow a similar model to that from Ref. [27].

Due to potential mechanical misalignment of the sensors or lenses, the actual initial ray position may deviate from the nominal one:

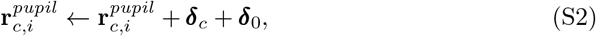

where ***δ***_*c*_ = (*δ*_*c,x*_, *δ*_*c,y*_, *δ*_*c,z*_) is the 3D translational misalignment of the *c*^*th*^ camera and ***δ***_0_ = (*δ*_0,*x*_, *δ*_0,*y*_, *δ*_0,*z*_) is a global 3D translational misalignment of the whole array to simplify optimization. Similarly, the whole array can be tilted, leading to a global optical axis tilt:

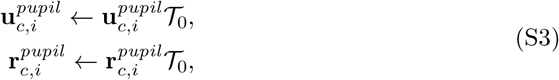

where *T*_0_ is a 3D rotation matrix, accounting for roll, pitch, and yaw. In other words, ***δ***_*c*_ + ***δ***_0_ and *T*_0_ constitute the 6-degree-of-freedom camera pose error.

These per-camera, per-pixel rays are propagated to the parabolic mirror by computing their distances from the misaligned pupil plane to the mirror surface,

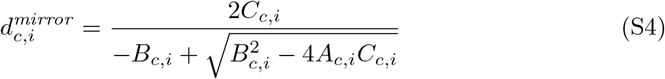

where

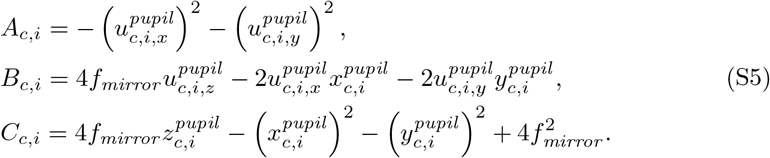

The position of each ray at the mirror surface is thus

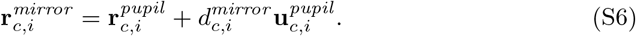

Upon reflection, the new ray direction is

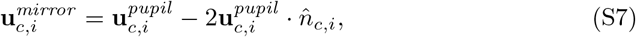

where 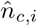 is the surface normal unit vector, whose unnormalized form is given by

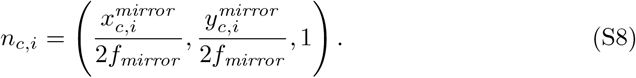

**Fig. S1.**
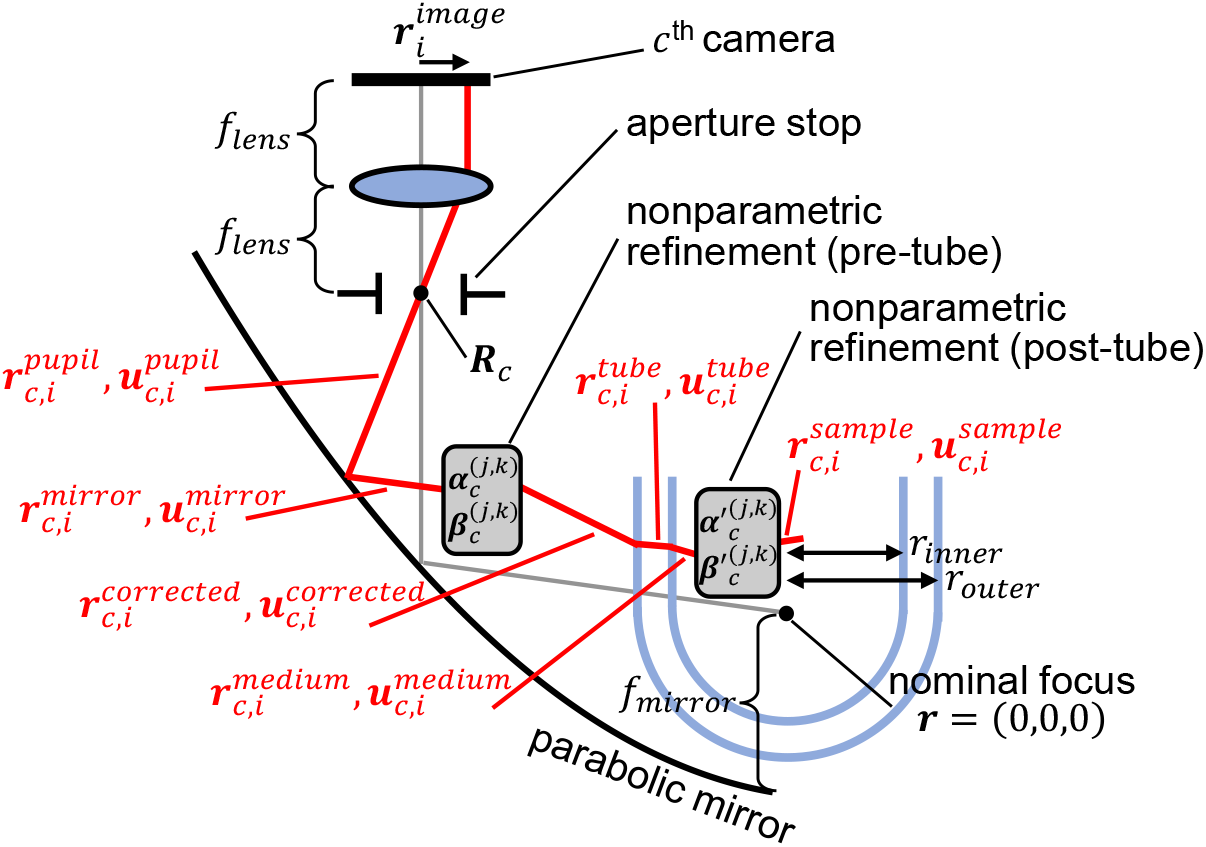
Ray propagation through the ReFLeCT system. A sample chief ray from the *ith* pixel of the *cth* camera is propagated (red). See main text (Sec. 2.3) and the supplement (Sec. S1) for further explanations, equations, and symbol definitions. Figure not drawn to scale.

Finally, we propagate the rays from the mirror surface to the region near the parabolic mirror focus:

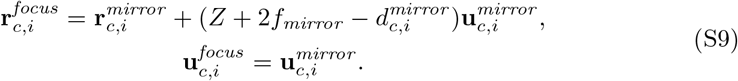

Note that the propagation distance is adaptive and defined so that every ray travels the same distance overall, and that rays parallel to the mirror’s optical axis always are reflected to the mirror’s focus.

#### S1.2 Accounting for misalignment and distortions (not from sample holder)

Apart from relative pose differences between the camera array and the parabolic mirror, the ray propagation outlined in Supplementary Sec. S1.1 assumes ideal conditions. In practice, the lenses may exhibit unpredictable distortions and misalignments and the mirror itself may not be perfectly parabolic, for example. To account for such distortions and misalignments, we do a nonparametric refinement of the rays in Eq. S9 using order-4 polynomials in the nominal lateral coordinates:

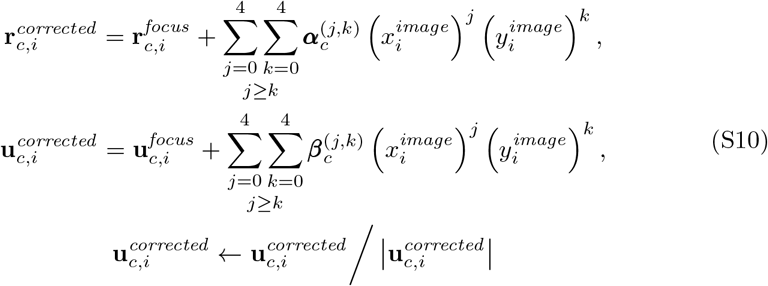

where 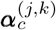 and 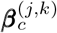 are both independently optimizable polynomial coefficients.

#### S1.3 Ray propagation through the sample holder

So far, we have only propagated rays in free-space. Next, we propagate the rays through the water-filled NMR tube, which consists of a hemispheric bottom that transitions into a cylindrical shaft. Thus, the NMR tube can be described by its intrinsic parameters: inner radius *r*_*inner*_, outer radius *r*_*outer*_, wall refractive index *n*_*glass*_, and medium refractive index *n*_*medium*_. In addition, we also model the tube’s extrinsic parameters, namely its 3D translational misalignment Δ**r**_*tube*_ and its 3D rotational misalignment, described by the axis-angle representation, 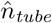 and *θ*_*tube*_ to form a 3D rotation matrix, *T*_*tube*_ (technically, due to the rotational symmetry of the NMR tube, the axis-angle representation has one redundant parameter).

First, to account for the tube pose change, we simply translate and rotate the rays, 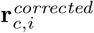 and 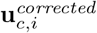, from Eq. S10:

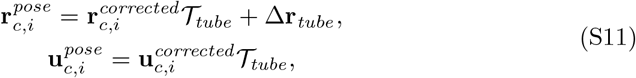

so that the center of the hemispherical part of the tube is nominally at the origin of the coordinate system. This simplifies the calculations for the subsequent ray propagation steps.

Next, we propagate the rays to the outer tube surface. To do this, we have to handle two possible intersections – with the spherical surface and the cylindrical surface. For the spherical case, the position of intersection is

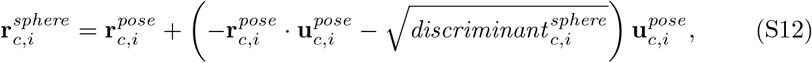

where

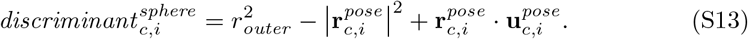

For the cylindrical case, the position of intersection is

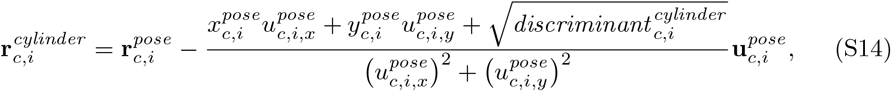

where

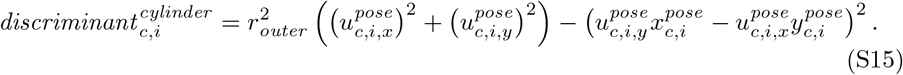

To determine whether to use the spherical or the cylindrical solution, we check which of the discriminants (Eqs. S13 and S15) is non-zero on a per-ray basis (i.e., per (*c, i*)), and assigned the corresponding solution to 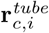 We then refracted the rays at the corresponding surface according to Snell’s law,

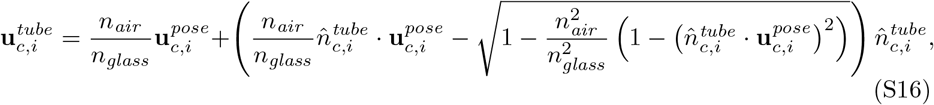

where 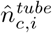 depends on whether we’re refracting at the spherical surface,

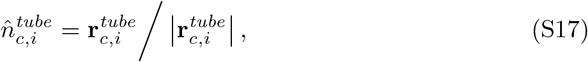

or the cylindrical surface,

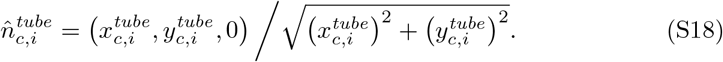

We then repeat Eqs. S12-S18 to propagate the rays to and refract at the inner surface, using different refractive indices and *r*_*inner*_ instead of *r*_*outer*_, to obtain 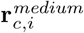 and 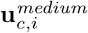. Finally, we are able to propagate through the medium in which the sample is freely moving.

In practice, the NMR tube may have manufacturing imperfections causing deviation from sphericity and cylindricity, which we account for by incorporating another nonparametric refinement, identical to the model described in Supplementary Sec. S1.2, except with a polynomial order of 15, with coefficients 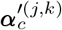 and 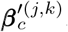 After this second nonparametric refinement, we obtain the final ray trajectories, 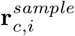 and 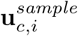

In the next two sections (Secs. S2 and S3), we describe how to optimize these parametric and nonparametric calibration parameters.

### S2 Camera array and parabolic mirror calibration with a translating fiber optic diffuser

We developed a joint hardware-software method for calibrating the misalignments and imperfections of both the parabolic mirror and the camera array. The hardware component consists of a fiber optic cannula with a diffuser tip with a 200-µm core diameter (Thorlabs), mounted on 3-axis motorized translation stage (Zaber) (Fig. S2). The diffuser at the tip of the cannula fiber scatters light in all directions, serving as a pseudo point source, so that it is visible from view points beyond 2*π* steradians. We scan the fiber tip in a known 3D pattern (i.e., a 9 × 9 × 9 grid, spanning 5 mm × 5 mm × 5 mm) across the 3D FOV of our imaging system and capture snapshots across all cameras for each scan position. The fiber tip shows up as a single small point in each camera image, allowing for straightforward segmentation and localization as feature points for 3D registration to calibrate the system misalignments and imperfections. Owing to the fact that there is physically only a single sample point at a given time, feature matching is trivial.

We developed a backwards bundle adjustment (BA) algorithm that allows us to use the sensor-to-sample ray propagation (described in Sec. 2.3 and Supplementary Sec. S1), which is reversed with respect to traditional BA algorithms. In other words, our backwards BA algorithm computes the *backprojection error* in the sample space rather than the *reprojection error* in the camera space in traditional BA. We do this by minimizing the shortest distance between each ray and the object point to which it corresponds. That is, given an object point with known 3D coordinates for the *k*^*th*^ stage scan position 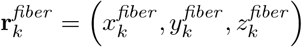, and the pixel *i*(*k*) in camera *c* that observes that *k*^*th*^ point, whose propagated ray is given by 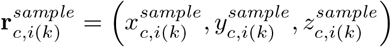 and 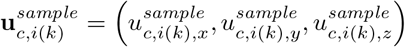, the shortest distance is given by

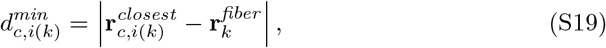

**Fig. S2.**
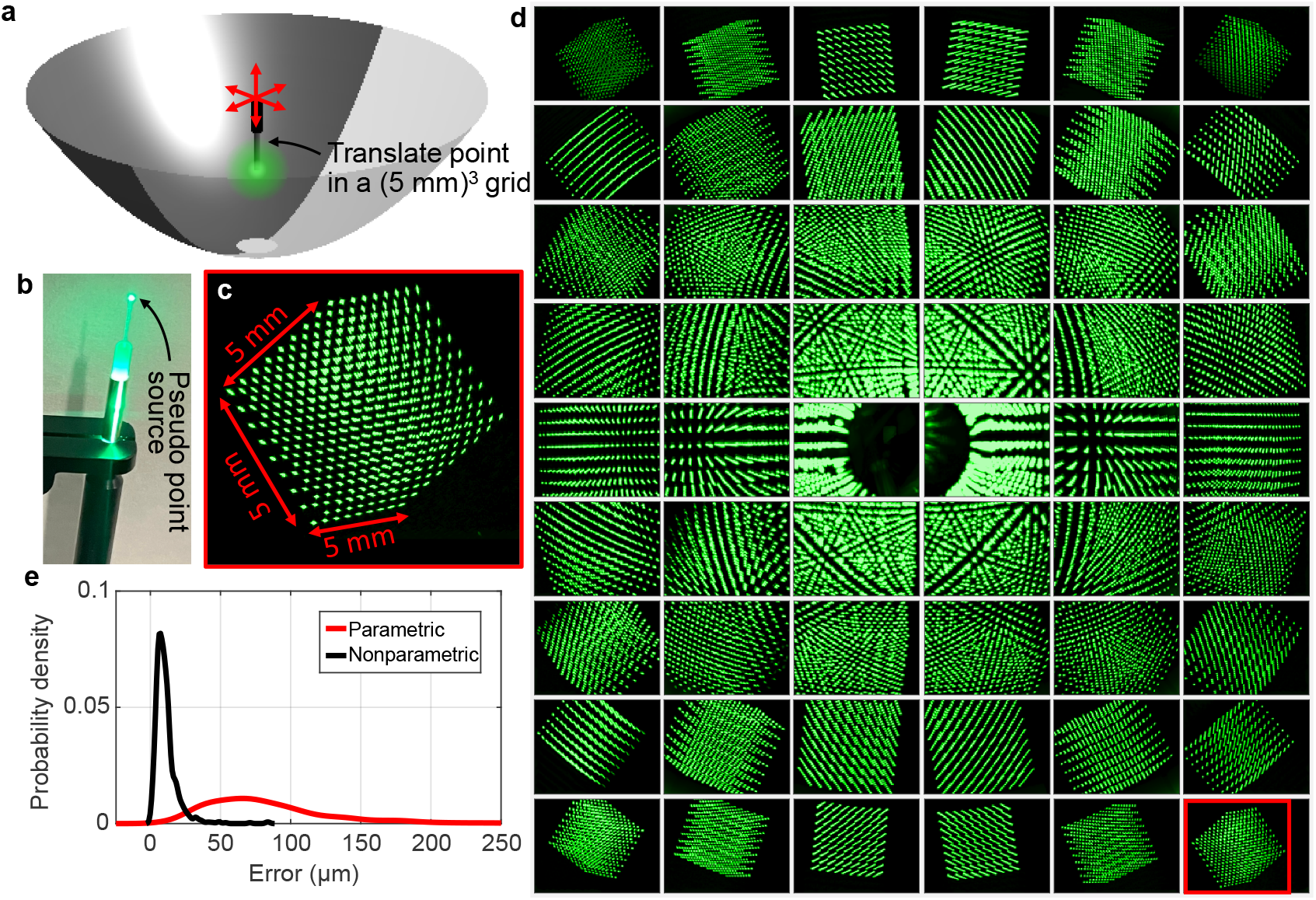
Calibration of the camera array and parabolic mirror. **b**, A point source is translated in a known 3D grid. **b**, Photo of the fiber optic cannula with a diffuser tip, serving as a pseudo point source visible from all directions. **c**, A single view of the 3D grid, formed by the fiber tip translation. **d**, Sample calibration dataset, consisting of multi-view data of the 3D grid. **e**, Distribution of distances between the fiber tip locations (estimated using the backwards bundle adjustment algorithm; Supplementary Sec. S2) and the ground truths, using the parametric model (assuming tilting and translation of perfect optics) and the nonparametric model (additionally using high-order polynomials).

where

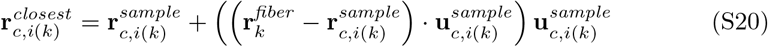

is the closest approach of the ray to the *k*^*th*^ fiber tip position. Thus, we minimize the mean square distance of every ray to its corresponding 3D fiber tip position

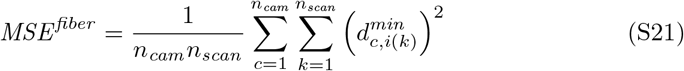

with respect to the optical system calibration parameters using gradient descent, where *n*_*cam*_ = 54 is the number of cameras and *n*_*scan*_ = 9^3^ = 729 is the number of fiber tip scan positions. Although in principle the ground truth object point locations are known, in practice the known trajectories may be off by a global 3D rotation and 3D translational offset, depending on the relative pose between the stage and the optical system. Furthermore, if the three translational axes are not perfectly orthogonal to each other, there may also be shear. Thus, in addition to optimizing the optical system parameters, we also jointly optimize an additional 8 parameters independent of *k*, describing 3D translation (Δ**r**), 3D rotation (*𝒯*), and 2D shear (*𝒮*),

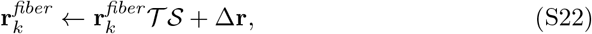

which is substituted into Eqs. S19 and S20.

Our joint hardware-software calibration approach is advantageous over a random sparse distribution of point emitters (e.g., fluorescence particles) for a few reasons. First, since the fiber tip is in air, we do not need to model the sample holder’s optics at the same time, thus simplifying the calibration problem. Second, our approach does not require matching points across different images, which may be thwarted by similarity in appearance of different point emitters within the sample. Third, since the trajectory of the fiber tip is pre-programmed, we know the ground truth locations (to within the stage accuracy).

#### S2.1 Two-step backwards bundle adjustment optimization

We perform gradient-descent-based minimization of Eq. S21 in two steps. First, we minimize *MSE* ^*fiber*^ with respect to the physically interpretable parameters: a common lens focal length *f*_*lens*_, parabolic mirror focal length *f*_*mirror*_, the relative pose between the camera array and mirror specified by ***δ***_0_ (Eq. S2) and *𝒯*_0_ (Eq. S3), and the intertranslation-stage calibration errors, specified by Δ**r**, *𝒯*, and *S* (Eq. S22). This first step obtains a close result, but still has significant errors (Fig. S2e). Thus, we perform a second gradient descent minimization of *MSE* ^*fiber*^, this time with respect to Δ**r**, *𝒯*, and *S* (Eq. S22) as well as the polynomial model parameters 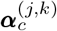 and 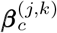 (Eq.S10). This results in a much better registration (Fig. S2e).

#### S2.2 Backwards bundle adjustment in the case of unknown calibration object positions

For the sake of completeness, we also consider the case for backwards BA where the object points are unknown. Here, the object points can be treated as optimizable variables or computed as the points that minimizes the closest distance squared to all rays that correspond to the same object point (hence, feature-matching is still required). That is, given a collection of rays, 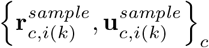, corresponding to unknown fiber (or fluorescent bead) *k*, the “best” point of intersection can be shown to be

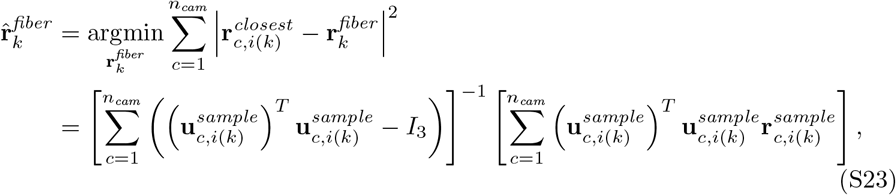

where *I*_3_ is the 3 *×*3 identity matrix and ^*T*^ is the transpose operation (here, creating column vectors from row vectors). We can then substitute 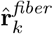 into Eqs. S19-S21 and proceed in the same way.

### S3 Sample holder calibration

Once the lenses and parabolic mirror are calibrated, we then need to calibrate the distortions stemming from the arbitrary pose and imperfections of the sample holder (the glass NMR tube). This NMR tube calibration is done in the following steps.

#### S3.1 Coarse estimation of NMR tube pose based on sample motion

Once the free-space calibration is complete, we begin coarse estimation of the NMR tube pose. Here, we rely on the sample motion within the lumen of the tube across many video frames. In particular, we take a max projection across entire videos or multiple videos and threshold the result, so that the resulting value for the *i*^*th*^ pixel of the *c*^*th*^ camera is

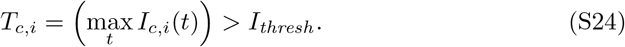

Thus, if the organism thoroughly explores the tube, then the thresholded max projection would be 1 inside the tube lumen and 0 outside. The goal for this coarse calibration step is to optimize the tube pose parameters so that the tube encloses the thresholded pixels (Eq. S24). To do achieve this, we specify a registration loss from the discriminants that were obtained from intersecting the rays with the tube surface (Eqs. S13 and S15). The key idea is that the discriminant is positive for rays that intersect with the tube surface and negative for rays that do not. Thus, we can form a prediction of 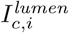 by sandwiching the discriminant between 0 and 1 using a logistic function,

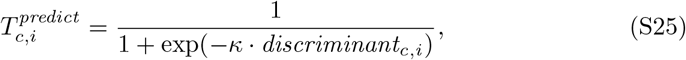

where *κ* is a scalar constant that tunes the steepness of the transition at the tube boundary to allow gradients to propagate through the discriminant. We thus minimize a weighted MSE,

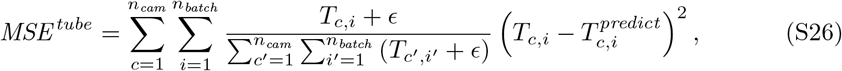

with respect to the tube pose parameters, 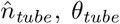, *θ*_*tube*_, and Δ**r**_*tube*_ (Eq. S11). Here, *n*_*cam*_ = 54 is the total number of cameras and *n*_*batch*_ is the number of pixels per camera in the gradient descent minibatch. The normalized weight factor in Eq. S26 confers greater importance to within-tube pixels, where *ϵ* tunes that importance – when *ϵ* = 0, the pixels outside of the tube do not contribute to the error, while when *ϵ* is very large, all pixels contribute approximately equally. The purpose of this weighting scheme is to account for situations when the organism doesn’t fully explore the tube lumen, by erring on the side of capturing more within-tube pixels inside the registered tube (i.e., prioritizing increasing the true positive rate over reducing the false positive rate).

#### S3.2 Refining tube pose jointly with a low-resolution sample reconstruction

We then refine the tube pose parameters jointly with reconstructing the moving sample at multiple time points at low resolution (10*×*-20*×* downsampled in all 3 spatial dimensions).

#### S3.3 Nonparametric refinement of sample-incident rays jointly with a low-resolution sample reconstruction

Finally, we update the polynomial model, with coefficients 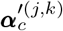 and 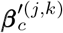 (mentioned at the end of Supplementary Sec. S1.3). This nonparametric model accounts for any residual deviations in the sample-incident ray trajectories, which stem from imperfections in the tube geometry and any remaining accumulated errors from earlier calibration steps. These polynomial coefficients are jointly updated with low-resolution (5 *×*downsampled in all 3 spatial dimensions) reconstructions of the sample at multiple time points.

After this second nonparametric refinement (the first one described in Supplementary Sec. S2), we obtain the final ray trajectories, 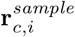 and 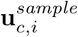.

### S4 Occupancy grid

After fully calibrating the ray trajectories, but before actually performing the dynamic volumetric reconstructions (Sec. 5.3), we generate occupancy grids for each video of interest. This procedure involves performing a low-resolution (12.5 *×*downsampled) 3D reconstruction for each time point and thresholding the fluorescence channels of the results. We then use 3D binary image dilation on the thresholded images to err on the side of having false positive voxels, even at the expense of false negative pixels to ensure no parts of the organism are erased. During high-resolution 3D reconstructions, each point along the ray traces are rounded to the nearest voxel in the occupancy grid. As the occupancy grids are binary (1 bit), they take up very little storage space and memory.

### S5 Supplementary videos

1. **Overview of ReFLeCT**.
2. **4D ReFLeCT imaging of a *Drosophila* larva with GFP-labeled myosin at 120 vps**.
3. **4D ReFLeCT imaging of a *Drosophila* larva expressing GFP in its pericardial cells at 30 vps**.
4. **4D ReFLeCT imaging of two *Drosophila* larvae expressing GFP in their salivary glands at 30 vps**.
5. **4D ReFLeCT imaging of two GFP-expressing zebrafish larvae at 120 vps**.
6. **4D ReFLeCT imaging of a GFP-expressing zebrafish larva at 120 vps**.
7. **4D ReFLeCT imaging of a zebrafish larva expressing GFP in its heart at 30 vps**.

### S6 Additional figures

**Fig. S3.**
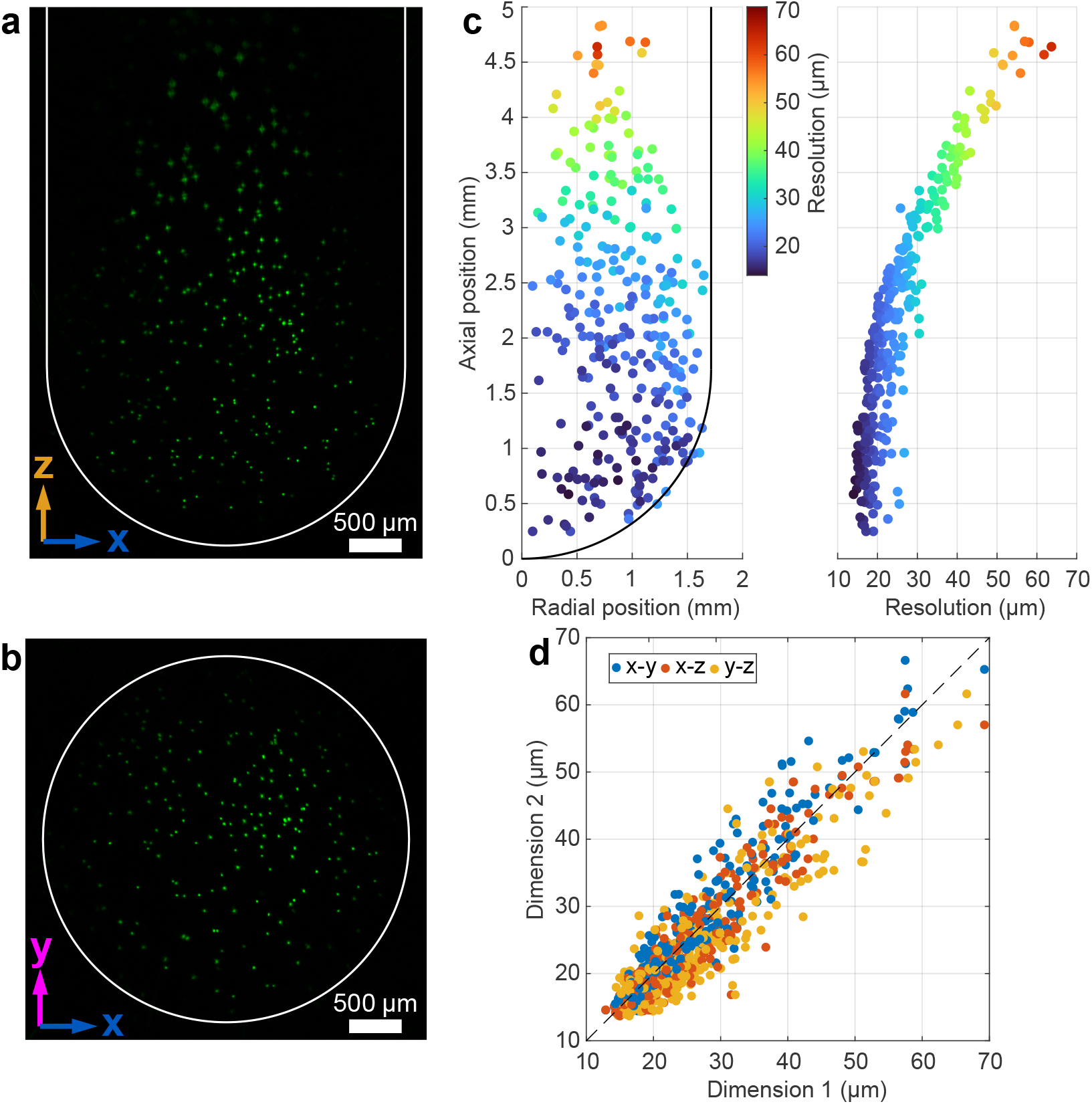
Resolution and FOV characterization of ReFLeCT with 6-*μ*m green fluorescent beads embedded in 1% agarose. **a**, *xz* projection of volumetric reconstruction. White line indicates the approximate inner boundary of the NMR tube. **b**, *xy* projection of the reconstruction across the bottom 2.4 mm of the tube. White circle indicates the approximate inner boundary of straight part of the NMR tube. **c**, FWHM of the beads, based on 3D Gaussian fits, of the fluorescent beads plotted across radial and axial position within the tube. FWHM values are the geometric mean values of the widths in the 3 dimensions. **d**, The *x, y*, and *z* FWHM values plotted pairwise against each other, indicating nearly isotropic resolution.

**Fig. S4.**
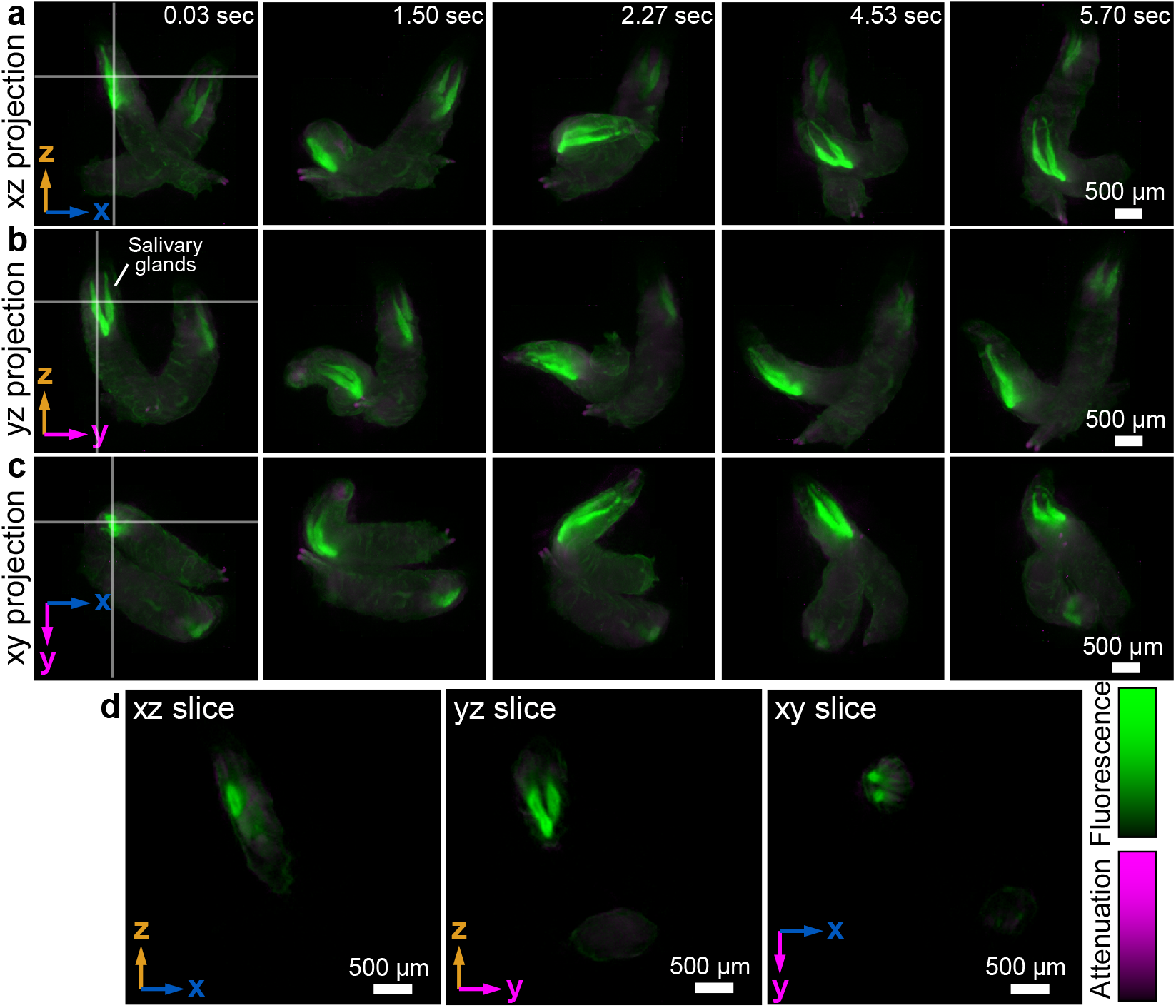
Multi-organism volumetric imaging. Two WL3 fruit fly larvae (*NP5169 Gal4>UAS-GFP-NLS*) expressing GFP in their salivary glands, imaged at 30 vps. See also the associated Video 4. **a**-**c**, Max intensity projections across all three imaging-system-centric axes, plotted at several time points. **d**, Cross-sections of the 3D reconstruction at 0.03 sec, denoted as vertical and horizontal lines in **a**-**c**. Note that our method reconstructs the hollow tube shape of the glands.

